# Basal gland localization and focal distribution of OLFM4-expressing cells in increasing severity of gastric intestinal metaplasia

**DOI:** 10.64898/2026.05.14.725297

**Authors:** Anuja Sathe, Rithika Meka, Benedikt Geier, Renee Long, Charlotte Wong, Ignacio A Wichmann, Summer Han, Jeanne Shen, Manuel R. Amieva, Hanlee P. Ji, Robert J. Huang

**Affiliations:** Division of Oncology, Department of Medicine, Stanford University School of Medicine, Stanford, CA, USA; Department of Microbiology and Immunology, Stanford University, Stanford, CA, USA; Department of Pediatrics, Stanford University, Stanford, CA, 94305; Division of Gastroenterology and Hepatology, Department of Medicine, Stanford School of Medicine, Stanford, CA, USA; Quantitative Sciences Unit, Department of Medicine, Stanford University School of Medicine, Stanford, CA, USA; Department of Neurosurgery, Stanford University School of Medicine, Stanford, CA, USA; Department of Pathology, Stanford University, Stanford, CA, USA

## Abstract

Patients with gastric intestinal metaplasia (**GIM**), a precancerous lesion, are at high risk for progressing to gastric cancer. Identifying these patients is critical to enable gastric cancer interception. Current approaches rely primarily on histologic evaluation of GIM severity and extent, which may be improved by incorporating molecular features that distinguish high-risk lesions. Our prior single-cell and spatial transcriptomics study identified differentially expressed genes associated with the highest-risk category of GIM. They included *ANPEP* expressed in enterocytes and *CPS1* and *OLFM4* expressed in intestinal stem-like or progenitor cells. We evaluated the protein expression and localization of these three markers to understand the cellular features associated with GIM risk and their spatial distribution within metaplastic tissues.

Using multiplex immunofluorescence, whole slide image analysis and confocal microscopy, we examined protein expression from 100 tissue biopsies annotated for metaplasia severity using the Operative Link on Gastric Intestinal Metaplasia Assessment (**OLGIM**) system. Tissue samples included control gastric tissue, GIM, dysplasia and adenocarcinoma. Quantitative whole slide image analysis demonstrated that CPS1 expression had a modest association with disease severity. Although ANPEP was strongly associated with GIM severity, it was also frequently expressed in stromal regions outside epithelial glands. In contrast, OLFM4 expression was largely restricted to epithelial glands and showed a strong association with increased OLGIM severity. These OLFM4-positive epithelial cells were present in discrete glandular foci that expanded with increasing severity of metaplasia. Within individual metaplastic glands, OLFM4 expression was highest at the gland base with decreased expression toward the gland surface.

Overall, these findings identified OLFM4 as a protein marker associated with high-risk GIM. The spatial organization of OLFM4-expressing cells at the base of metaplastic glands and their focal expansion within tissues suggest the presence of a stem cell-like epithelial compartment that may contribute to the progression of GIM towards gastric cancer.

## INTRODUCTION

Gastric cancer is the fifth most common cancer worldwide and remains a leading cause of cancer-related mortality^1^. Gastric adenocarcinoma of the intestinal subtype, representing the majority of cases, arises through a multistep process of atrophy, metaplasia and dysplasia, commonly referred to as the Correa cascade^2^. Within this process, gastric intestinal metaplasia (**GIM**) represents a key precursor lesion associated with increased cancer risk and provides an opportunity for early gastric cancer interception.

Although GIM is a precancerous lesion, only a subset (approximately 3-5%) of patients with GIM progress to gastric cancer^3–7^. Improving surveillance and interception strategies requires identifying patients that are at high risk of cancer progression. Currently the most robust clinical risk stratification algorithm is the Operative Link on Gastric Intestinal Metaplasia Assessment (**OLGIM**) system, which grades the severity of GIM using both the extent of glands involved at the microscopic level, and the anatomic extent of disease^8^. Higher OLGIM stages are consistently associated with increased risk for gastric cancer^9,10^. However, OLGIM staging is based on morphologic and histopathology assessment alone - OLGIM does not capture molecular features that may underlie disease progression. Moreover, OLGIM demonstrates limited discrimination in predicting the subset of individuals who progress on to neoplasia. In risk prediction models, this corresponds to only moderate predictive performance, with reported AUROC/C-statistic values of approximately 0.7^11^. Incorporating novel biomarkers has the potential to improve the identification of patients at high-risk of progression to GC^12^.

We previously identified a 26-gene signature that was associated with advanced OLGIM stages and localized to metaplastic glands on histopathology^13^. Based on gene expression analysis of patient biopsies, this signature was significantly expressed in mature and immature intestinal cells not normally present in the healthy stomach. This result indicated molecular reprogramming within metaplastic epithelium associated with an increased risk of progression. While this study identified novel gene expression markers associated with high-risk GIM, it remains unresolved if these markers are also upregulated at the protein level. A gene expression signature is more difficult to implement for widespread clinical application. Ideally, any risk-prediction molecular assay would be compatible with standard pathology workflows across diverse settings (such as smaller community hospitals). Finally, the spatial organization of these markers within metaplastic glands remains poorly understood.

To address these limitations, we evaluated the protein expression of several important markers, of the high-risk gene expression signature, namely ANPEP, OLFM4 and CPS1. On GIM biopsies, we used tyramide signal amplification based multiplex immunofluorescence (**mIF**) staining for these proteins. ANPEP (aminopeptidase N, CD13) is a membrane-bound metalloprotease involved in peptide digestion at the brush border of intestinal epithelial cells^14^. CPS1 (carbamoyl phosphate synthetase 1) is a mitochondrial enzyme that catalyzes the rate-limiting step of the urea cycle^15^. OLFM4 (olfactomedin-4) is a secreted glycoprotein with roles in mucosal differentiation, inflammation and defense against pathogens^16^. In our previous study^13^, single-cell RNA sequencing (**scRNA-seq**) analysis demonstrated that these genes were expressed by distinct epithelial lineages in GIM. *ANPEP* was expressed by mature enterocytes, while *OLFM4* and *CPS1* were expressed by immature intestinal lineages such as stem or progenitor cells. Similar cellular patterns of expression have also been identified by other scRNA-seq or spatial transcriptomics studies of GIM^12,17,18^. Hence, this three-marker mIF panel enabled the simultaneous evaluation of both molecular and cellular features associated with GIM risk. Using whole slide image (**WSI**) analysis and confocal microscopy, we examined the cellular expression patterns of ANPEP, CPS1 and OLFM4 in GIM tissues and quantified their association with GIM severity in gastric tissue derived from a cohort of patients with histopathology spanning the neoplastic spectrum. In addition, we examined their spatial organization within metaplastic tissues and glands to understand the cellular architecture of GIM and early neoplasia.

## RESULTS

### Patient cohort and study design

Our study included 100 biopsies from 94 patients who underwent upper gastrointestinal endoscopy (**Table 1**). Each biopsy was graded for the severity of intestinal metaplasia using the Sydney system visual analog scale as part of OLGIM staging by a dedicated gastrointestinal pathologist (JS)^19^. Biopsies underwent histopathologic review and were scored as mild GIM, moderate GIM or marked GIM (**Table 2**). Controls included biopsies with normal mucosa (‘normal’), and those with normal mucosa from patients with GIM identified in additional biopsies (‘GIM-adjacent’). The human stomach is a complex organ with tissue-level differences based on location (*e.g.* antrum vs corpus). As most carcinomas occur in the distal stomach^20^ and to control for subsite differences, all GIM and control biopsies were obtained from the gastric antrum. In addition, we included biopsies with dysplasia or cancer from either the antrum or the incisura.

**Table 1:** Clinical characteristics of the patient cohort.

| Variable | N | Percent |
| --- | --- | --- |
| **Age (years)**, mean ± standard deviation | 63.7 ± 11.4 | |
| **Age group** |  |  |
| <50 years | 11 | 11.7 |
| 50–59 years | 19 | 20.2 |
| 60–69 years | 35 | 37.2 |
| $\geq 70$ years | 29 | 30.9 |
| **Sex** |  |  |
| Female | 56 | 59.6 |
| Male | 38 | 40.4 |
| **Race** |  |  |
| Asian | 52 | 55.3 |
| Black | 1 | 1.1 |
| Other | 3 | 3.2 |
| Unknown | 19 | 20.2 |
| White | 19 | 20.2 |
| **Ethnicity** |  |  |
| Hispanic/Latino | 18 | 19.1 |
| Non-Hispanic | 75 | 79.8 |
| Unknown | 1 | 1.1 |
| **OLGIM stage** |  |  |
| 0 | 19 | 20.2 |
| 1 | 22 | 23.4 |
| 2 | 20 | 21.3 |
| 3 | 20 | 21.3 |
| 4 | 13 | 13.8 |
| **Patients with more than one biopsy** | 6 | 6.4 |

**Table 2:** Sample characteristics for GIM severity determined by histopathology.

| Variable | % |
| --- | --- |
| Normal | 18 |
| GIM-adjacent | 6 |
| GIM-mild | 20 |
| GIM-moderate | 20 |
| GIM-marked | 27 |
| Dysplasia | 4 |
| Cancer | 5 |

All biopsies underwent staining for *Helicobacter pylori* at time of collection and were negative. This was a control to limit the confounding effects of active *Helicobacter pylori* infection. We performed mIF on adjacent tissue sections from these formalin-fixed paraffin-embedded (**FFPE**) biopsies (**Methods**) to evaluate the cellular and spatial expression patterns of ANPEP, OLFM4 and CPS1 proteins which were identified as markers of high-risk GIM in our prior study^13^.

### Cellular expression patterns of ANPEP, CPS1 and OLFM4 proteins

We evaluated the cellular localization patterns of ANPEP, CPS1 and OLFM4 proteins. We included small intestine and tonsil as control tissues with known expression patterns of these markers (**Methods**). We used two imaging approaches. WSI provided the marker distribution across the entire biopsy. We also performed high-resolution confocal laser scanning microscopy on control tissues and representative samples from our cohort to evaluate subcellular localization of markers.

In the small intestine, ANPEP expression was localized intracellularly and at the apical membrane of the brush border epithelium formed by enterocytes, and additionally secreted into the gland lumen (**Supplementary Fig. 1A**) as described previously^14,21^. CPS1 was localized in the cytoplasm. OLFM4 subcellular localization was predominantly cytoplasmic with some extracellular secreted protein. In the tonsil, ANPEP was absent in the germinal center but detected in surrounding cells (**Supplementary Fig. 1B**), consistent with previously described expression in immune and stromal cells^22^. Weak expression of CPS1 was observed in the germinal center of the tonsil while OLFM4 was absent. For both the small intestine and tonsil, these expression and localization patterns matched results reported in the Human Protein Atlas using immunohistochemistry^23,24^. This result validated our mIF staining approach.

Next, we assessed protein localization across our set of control, GIM, dysplasia and cancer samples (**Fig. 1**, **Supplementary Fig. 2**). In GIM, we observed similar localization patterns of ANPEP as in the small intestine with high concentration in the brush border microvillar membrane. However, ANPEP was also expressed in non-glandular regions containing stromal and immune cells, similar to our observations in the tonsil. Intracellular CPS1 expression was detected in the glands of GIM, dysplasia and cancer with little expression in controls. OLFM4 expression in GIM, dysplasia and cancer were cytoplasmic and secreted. It was largely specific to glandular regions and rare in controls, although OLFM4 expression has been noted in neutrophils in prior literature^25,26^. These findings confirmed that specific proteins from the high-risk GIM gene signature we previously identified^13^ were detectable among GIM tissues.

**Figure 1.**
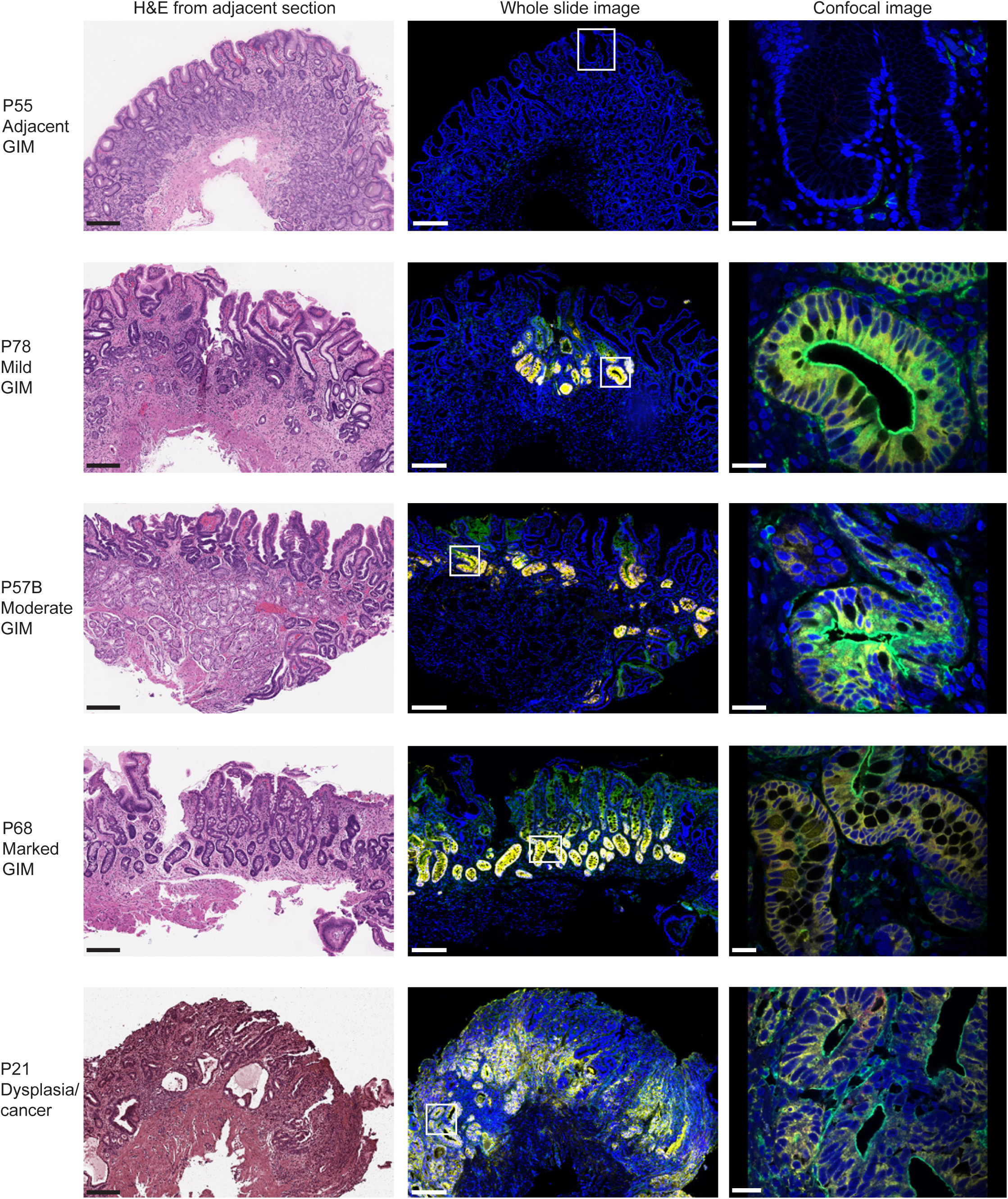
mIF staining of ANPEP, CPS1 and OLFM4 in gastric tissues. (A) H&E image from adjacent section (left panel), multiplex immunofluorescence whole slide image (middle panel) and confocal image (right panel) from respective samples. Rectangle with white box in the multiplex immunofluorescence whole slide image (middle panel) indicates area that was imaged for the confocal image (right panel). Fluorescence images are pseudocolored for DAPI (blue), ANPEP (green), CPS1 (red) and OLFM4 (yellow). Scale bars: 200 μm (left panel), 200 μm (middle panel), 20 μm (right panel).

### Whole-slide quantification of ANPEP, CPS1 and OLFM4 expression

We created a reproducible image analysis pipeline for mIF WSI using the QuPath software^27^ (**Methods**). Using watershed-based detection, we first performed cell segmentation (**Fig. 2A**). This identified 7.28 million cells from all images in the cohort with an average of over 72 thousand cells per image. Using a threshold-based classifier, we determined whether each cell was positive for ANPEP, CPS1 or OLFM4 protein expression (**Fig. 2B-D**, **Supplementary Fig 3A-C**). This approach enabled quantitative classification of protein expression status at the level of individual cells across all images.

**Figure 2.**
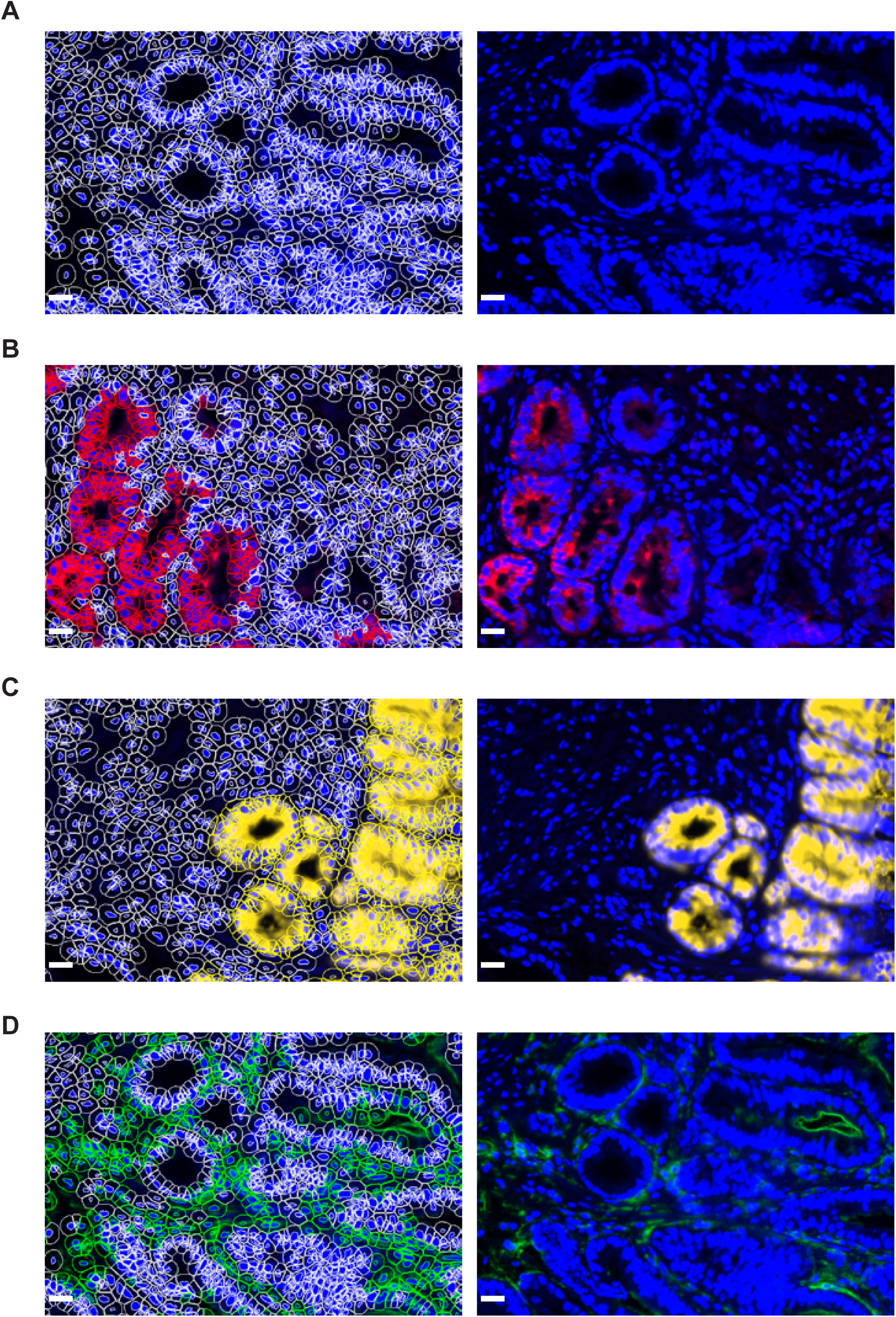
Whole slide image analysis using cell detection and classifiers. (A) Left panel: Overlay of outlines of detected cells and nuclei following cell segmentation algorithm in white with DAPI in blue. Right panel: DAPI in blue without overlay. (B) Left panel: Overlay of outlines of cells classified as positive for CPS1 expression (in red) or negative for CPS1 expression (white) together with CPS1 staining (red) and DAPI (blue). Right panel: CPS1 staining (red) and DAPI (blue) without overlay. (C) Left panel: Overlay of outlines of cells classified as positive for OLFM4 expression (in yellow) or negative for OLFM4 expression (white) together with OLFM4 staining (yellow) and DAPI (blue). Right panel: OLFM4 staining (yellow) and DAPI (blue) without overlay. (D) Left panel: Overlay of outlines of cells classified as positive for ANPEP expression (in green) or negative for ANPEP expression (white) together with ANPEP staining (green) and DAPI (blue). Right panel: ANPEP staining (green) and DAPI (blue) without overlay. Representative images from P34 are pseudocolored, scale bar 200 μm.

### OLFM4 expression is strongly associated with GIM severity

We next evaluated the association between protein expression and GIM severity. For each sample, we quantified the proportion of cells positive for ANPEP, CPS1 or OLFM4 identified using this classifier. The proportion of ANPEP-expressing cells was significantly higher in moderate and marked GIM compared to controls or mild GIM (FDR-adjusted *p* ≤ 0.05) (**Fig. 3A**). However, the baseline proportion of ANPEP-expressing cells in normal and adjacent GIM was already high (20.5 – 41.8%), likely reflecting its expression in stromal and immune cells beyond metaplastic epithelial cells. In addition, we tested this association using a linear regression model with GIM severity modelled using an ordinal scale (**Fig. 3B**). Proportion of ANPEP-expressing cells showed a significant positive association with stage (β = 4.54 percentage points per stage, R² = 0.19, *p* = 1.7×10⁻⁵).

**Figure 3:**
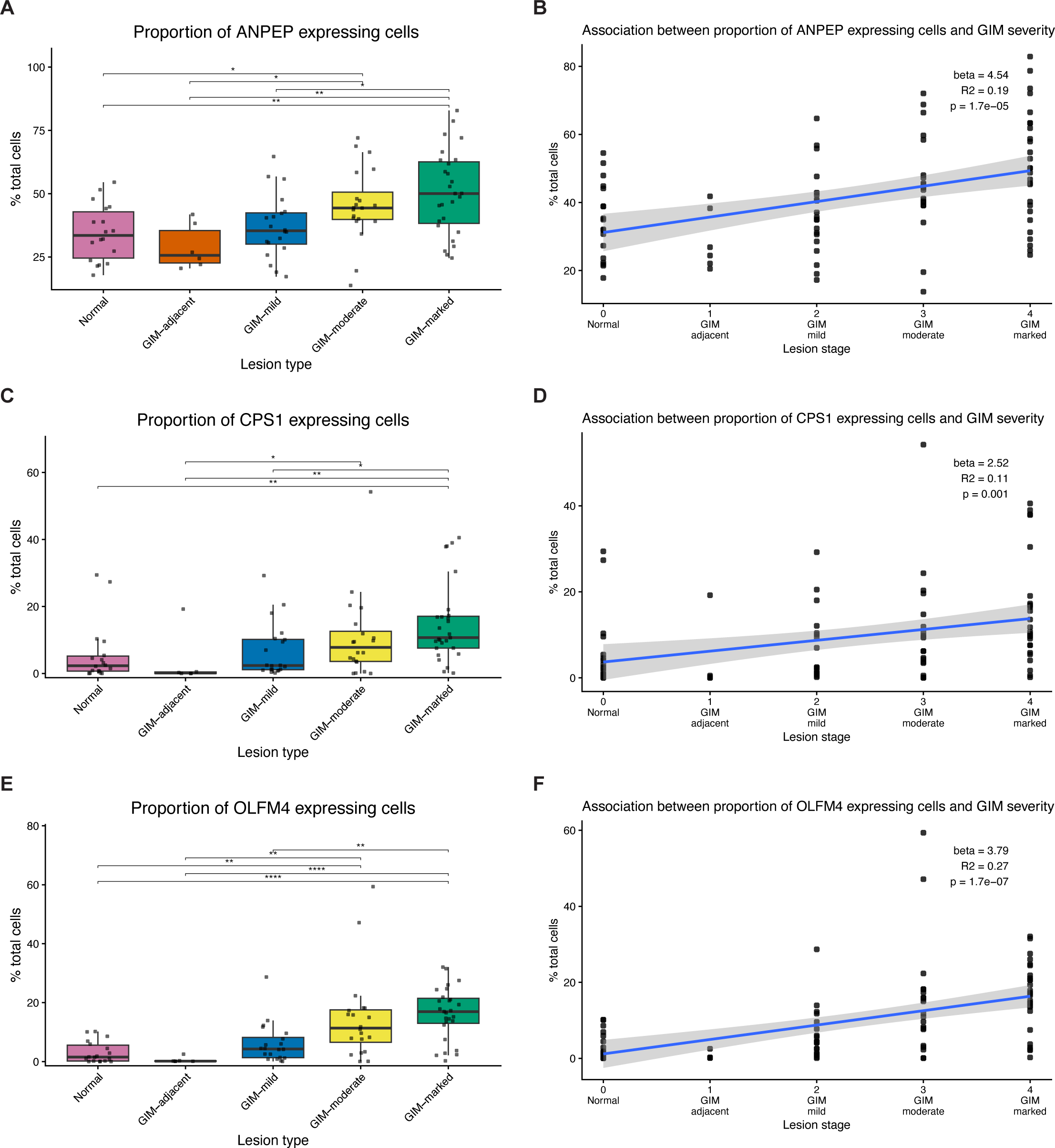
Association between proportion of cells expressing respective proteins and GIM severity. (A, C, E) Comparison of proportion of cells expressing the respective marker across lesion categories. Each point represents an individual sample, and boxplots show the median and interquartile range. Differences between groups were evaluated using the Kruskal–Wallis test followed by Dunn’s multiple comparisons test with false discovery rate correction. Only statistically significant comparisons are displayed in the figures. (B, D, F) Linear regression analysis testing the association between the proportion of marker-positive cells and lesion severity. Lesion categories were encoded as an ordinal stage variable. Each point represents one sample; the line indicates the fitted regression with 95% confidence interval, with regression coefficient (β), coefficient of determination (R²), and p-value.

CPS1 expression was low in normal and GIM-adjacent tissue (0–19.2%). The proportion of CPS1-positive cells was significantly higher in marked GIM compared to controls and mild GIM (FDR-adjusted *p* ≤ 0.05) (**Fig. 3C**). The linear regression modeling demonstrated a significant but more modest association with GIM severity (β = 2.52 percentage points per stage, R² = 0.11, p = 0.001) (**Fig. 3D**).

OLFM4 expression was minimal in controls and adjacent GIM (0.09-2.53%) and increased significantly in both moderate and marked GIM (FDR-adjusted *p* ≤ 0.05) (**Fig.3E**). Linear regression confirmed a strong positive association with severity (β = 3.79 percentage points per stage, R² = 0.27, p = 1.7×10⁻⁷) (**Fig. 3F**). While the expression of all three markers was associated with GIM, the proportion of OLFM4-expressing cells showed the most consistent increase with progression of GIM severity. Association of OLFM4 expression with severity was stronger than its colocalization with ANPEP (β = 2.33 percentage points per stage, R² = 0.18, p = 2.1×10⁻5), CPS1 (β = 1.69 percentage points per stage, R² = 0.17, p = 6.4×10⁻5 or both markers (β = 1.12 percentage points per stage, R² = 0.15, p = 1.6×10⁻4) (**Supplementary Fig. 3D-F**). Dysplasia and cancer samples maintained the expression of ANPEP, CPS1 and OLFM4 at levels comparable to marked GIM (**Supplementary Fig.3G**).

### OLFM4-expressing cells are present in discrete glandular foci in GIM

Next, we examined WSI images to understand the spatial distribution of these OLFM4-expressing cells within the gastric antrum. The distribution of OLFM4-expressing cells was localized only within certain areas of each tissue and not homogenously distributed (**Fig. 4**). While most normal and GIM-adjacent control samples did not have large numbers of OLFM4-positive cells, focal areas of OLFM4 expression were occasionally observed. In mild GIM, we observed discrete glandular foci of OLFM4-expressing cells. With increasing severity, these foci of OLFM4-positive cells occupied larger areas. These findings suggest that OLFM4-positive cells emerge in discrete small foci, first affecting individual glands and then expanding as clusters of affected glands with increased severity of GIM.

**Figure 4.**
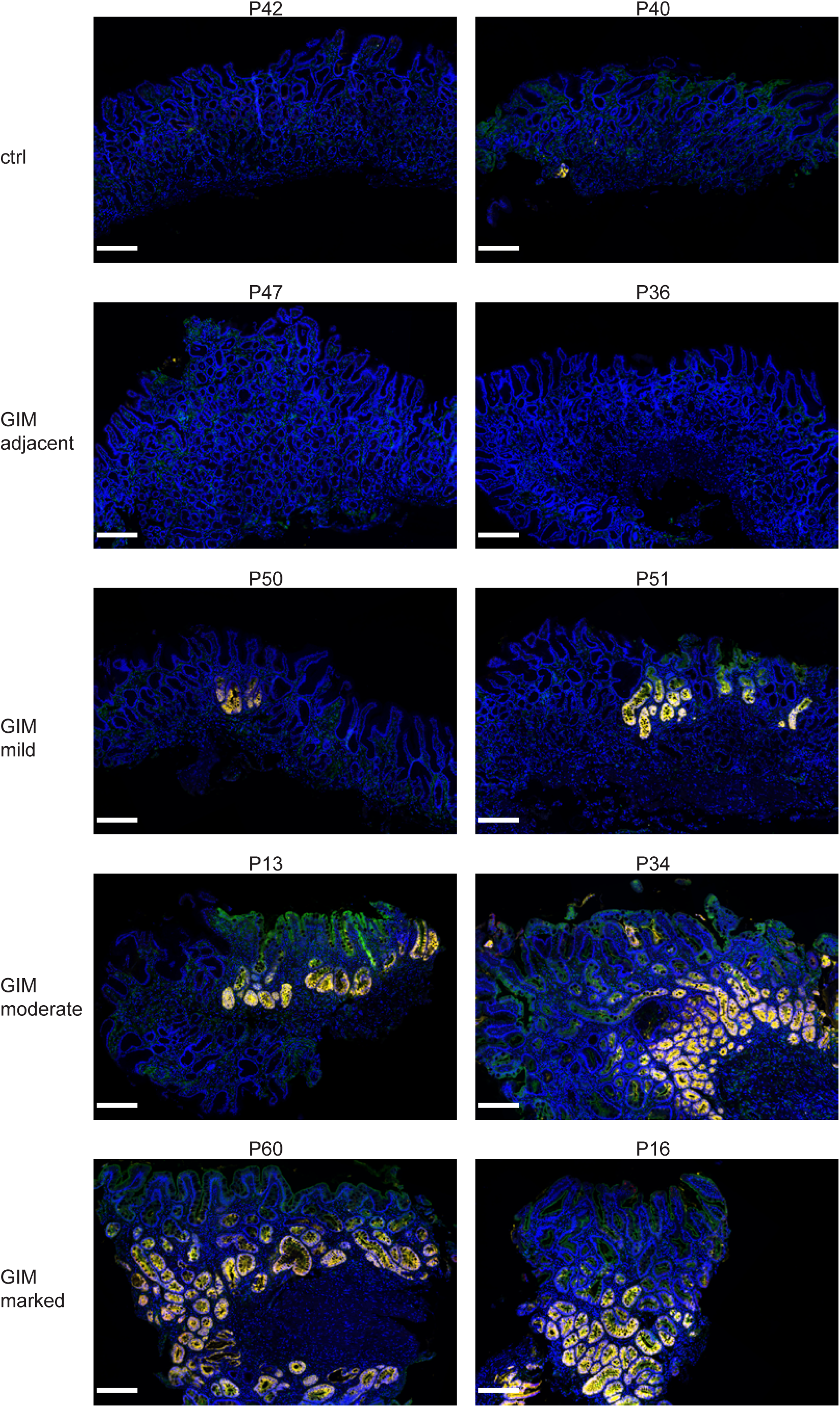
Tissue distribution of ANPEP, CPS1 and OLFM4 in gastric tissues. (A) Multiplex immunofluorescence whole slide images of respective samples across lesion categories. Fluorescence images are pseudocolored for DAPI (blue), ANPEP (green), CPS1 (red) and OLFM4 (yellow). Metaplasia marker expression is located to discrete glands, not globally. With increasing metaplastic severity, foci of OLFM4-expressing cells occupy larger areas. Scale bars: 200 μm.

### OLFM4-expressing cells localize to the base of metaplastic glands

Given that normal antral gland cells generally do not express OLFM4, we examined the spatial location of OLFM4-positive cells within metaplastic glands. For this analysis, we evaluated regions of interest from tissue samples where gland architecture was preserved along the longitudinal axis. Additionally, we evaluated their location in the small intestine where OLFM4-expressing stem cells are known to localize to the crypt base^28,29^.

In the intestine, we confirmed the restricted expression of OLFM4 in cells located at the crypt base (**Fig. 5A**). In GIM glands, OLFM4 had a distinct spatial pattern of expression (**Fig. 5B-C, Supplementary Fig. 4**). The highest expression was observed at the gland base, with fewer positive cells towards the gland surface. Unlike the intestine, where OLFM4 expression was confined to a relatively restricted crypt base compartment, a substantially larger population of gland base cells expressed OLFM4 in GIM. CPS1-expressing cells were distributed throughout the gland. ANPEP expression was highest at the enterocyte brush border near the gland surface but was also present along the longitudinal axis. Our results indicate that the cellular spatial architecture of GIM glands resembles intestinal glands, with a greater proportion of OLFM4-expressing cells localized to the gland base.

**Figure 5.**
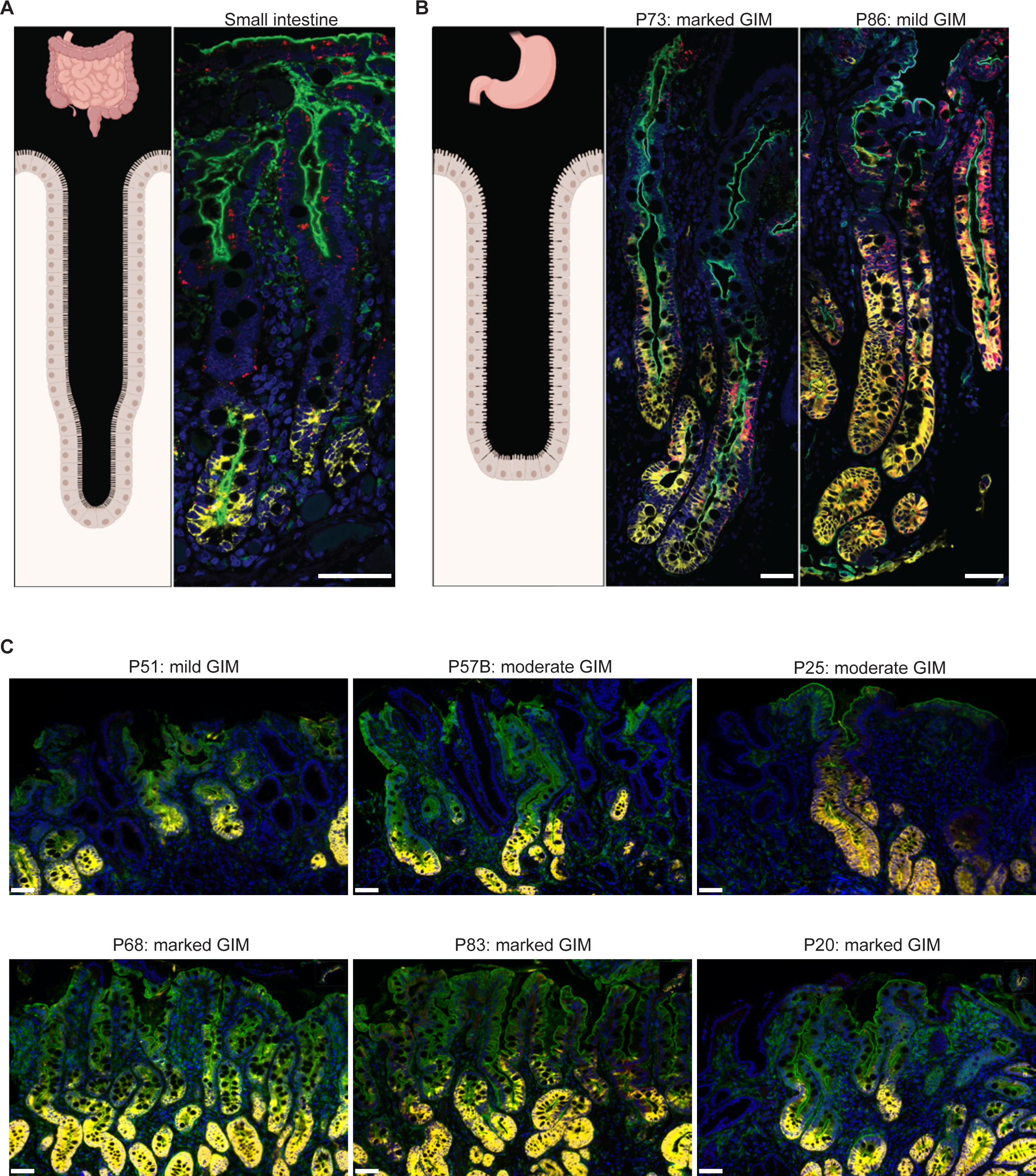
Spatial distribution of ANPEP, CPS1 and OLFM4. (A) Schematic representation of small intestine crypt (left panel) and confocal image of mIF staining (right). Scale bar: 50 μm. (B) Schematic representation of gastric antrum gland (left panel) and confocal images of mIF staining from respective samples (right). Scale bar: 50 μm. (C) Whole slide images from respective samples. Scale bar: 50 μm. All fluorescence images are pseudocolored for DAPI (blue), ANPEP (green), CPS1 (red) and OLFM4 (yellow).

## DISCUSSION

In this study, we used multiplex immunofluorescence combined with whole-slide image analysis and confocal microscopy to examine the protein expression and spatial organization of markers previously associated with high-risk GIM. Across a large cohort of gastric biopsies, we demonstrated that OLFM4-expressing epithelial cells emerge focally, expand with increasing GIM severity and exhibit a consistent basal localization within metaplastic glands.

Our findings extend our prior gene expression study that identified *ANPEP*, *CPS1* and *OLFM4* as markers of high-risk GIM^13^. We confirmed that these proteins are detectable in GIM tissues. OLFM4 expression demonstrated a strong and consistent relationship with GIM severity. Although a recent study used RNA in situ hybridization to demonstrate an association between *OLFM4* gene expression, GIM and gastric cancer^30^, they did not report the abundance, cellular and spatial localization of OLFM4 protein^31^. Our study includes quantitative evaluation of OLFM4 protein expression across a large well-annotated clinical cohort. We identified notable features regarding the spatial distribution of these cells at both the tissue level and within individual metaplastic glands.

ANPEP expression was associated with increased GIM severity, consistent with the fact that OLGIM staging reflects the extent of intestinal-type epithelium observed on histopathology^19^ and that ANPEP is highly expressed by enterocytes^21^. However, our analysis also revealed substantial ANPEP expression in stromal regions outside epithelial glands, reflecting expression in non-epithelial cell populations. These findings limit its utility as a quantitative epithelial marker of GIM. CPS1 expression showed only a weak association with GIM severity as reported previously^32^, similarly limiting its application as a potential quantitative biomarker.

At the tissue level, OLFM4-expressing cells exhibited a focal distribution within GIM tissues. While most control samples lacked significant OLFM4 expression, occasional small foci of OLFM4-positive glands were observed even in tissues classified as normal or GIM-adjacent. With increasing GIM severity, these foci expanded to involve larger areas of the mucosa. This spatial pattern suggests that OLFM4-expressing epithelial populations may arise in discrete glandular foci and subsequently expand locally. These observations are compatible with prior genomic studies demonstrating a clonal origin of metaplastic glands^12,33,34^ and their propagation within the gastric mucosa through gland fission^33^.

Within individual metaplastic glands, OLFM4-expressing cells showed a consistent spatial organization within GIM. OLFM4 expression was highest at the gland base. This result is analogous to the spatial localization of OLFM4-positive stem cells located at the base of intestinal crypts^28,29^. Stem cell populations in gastric antral glands have been proposed to reside in the isthmus, neck, or gland base^35,36^. The basal enrichment of OLFM4-expressing cells suggests that intestinal lineage programs in GIM may establish a stem-like epithelial compartment at the antral gland base that gives rise to differentiated epithelial populations along the gland axis. The origin of OLFM4-expressing epithelial cells, which are not typically present in healthy antral glands, remains uncertain. One cause may be the reprogramming of pre-existing gastric progenitor cells located at the gland base by *Helicobacter pylori* infection and inflammation. This hypothesis is consistent with studies demonstrating Lgr5-expressing cells at the base of antral glands in mice that function as long-lived gastric stem cells^37,38^. Alternately, OLFM4-expressing cells might arise from the migration of reprogrammed stem cells located in the isthmus, in line with lineage tracing studies in mice proposing the isthmus as the principal stem cell compartment^39,40^. Another possibility is that differentiated epithelial cells located at the gland base undergo transdifferentiation, the proposed mechanism underlying pseudopyloric metaplasia in the corpus^41,42^. Future studies investigating the molecular mechanisms underlying epithelial reprogramming, together with lineage tracing studies in human-derived models, can potentially resolve this. Single-cell multiomic approaches integrating chromatin accessibility and transcriptional state may provide additional insight into the regulatory mechanisms underlying acquisition of *OLFM4* expression. Spatial transcriptomic and proteomic approaches examining larger numbers of features may further define the molecular phenotype of these basally located OLFM4-expressing cells through co-expression analysis.

The spatial organization of OLFM4-expressing cells also suggests that progression of GIM may involve expansion of a stem-like epithelial compartment rather than simply an increase in the proportion of differentiated intestinal cell types. In early lesions, OLFM4-expressing cells were confined to a limited number of glands or small tissue regions. As metaplasia severity progressed, both the number of OLFM4-positive glands and the size of OLFM4-positive regions increased. These expanded OLFM4-expressing epithelial populations may be the cell type that drives the development of dysplasia and gastric cancer. Recent scRNA-seq studies have identified *OLFM4* as a marker of intestinal stem-like or progenitor cells in GIM. Computational lineage trajectories suggest that *OLFM4*-positive cells may give rise to reprogrammed metaplastic epithelial populations and share transcriptional features with early gastric cancer cells^12,13,17,18,43–48^.

Our findings also have potential clinical implications. Current risk stratification for GIM relies primarily on histopathologic assessment that is integrated with approaches such as OLGIM staging. However, these pathology-based approaches do not incorporate underlying molecular alterations within metaplastic epithelium. Moreover, there exists high inter-observer variability in GIM detection and severity assessment between pathologists^49,50^. OLFM4-based immunostaining may complement H&E-based staining to enhance detection, an intriguing possibility raised by our study’s finding of small foci of OLFM4-stained glands in GIM-adjacent tissue which was adjudicated as ‘normal’ by even a specialized pathologist. A GIM-specific marker may also prove useful in developing automated detection and staging strategies, such as training computer vision models to recognize GIM and to stage GIM histologic severity. Early-stage deep learning models based on H&E demonstrate limited accuracy in GIM detection^51–53^, which may be substantially improved by disease-specific markers.

Several limitations should be considered when interpreting our study’s findings. Our analysis was cross-sectional and did not include longitudinal follow-up to determine whether marker expression directly predicted progression to gastric cancer. We also did not examine potential differences between complete and incomplete intestinal metaplasia, and in lesions located in the stomach body. This cohort was also negative for *Helicobacter pylori*. WSI quantification relied on nuclear staining with watershed cell segmentation, which may not fully capture true cellular membrane boundaries. Extracellular secreted signal may not be captured by cell-level measurements. Finally, assessment of gland-axis spatial patterns required evaluation of glands with preserved longitudinal architecture that is not always available because of oblique planes in tissue sectioning.

## MATERIALS AND METHODS

### Samples

This study was conducted in compliance with the Helsinki Declaration. All patients were enrolled according to study protocols approved by the Stanford University School of Medicine Institutional Review Board (IRB-45077 and IRB-44036). Written informed consent was obtained from all patients. Tissues were processed using formalin fixation and paraffin embedding as a part of routine clinical pathology workflow. Hematoxylin and eosin (**H&E**) sections were graded for the severity of intestinal metaplasia using the Sydney system visual analog scale as part of OLGIM staging by a dedicated gastrointestinal pathologist (JS). For our study, we obtained 5 μm adjacent sections from the same tissue block. These sections were used for additional H&E or mIF staining.

### Multiplex immunofluorescence

We performed mIF staining as described previously^54^. FFPE sections were deparaffinized using xylene and hydrated in a descending alcohol series. Antigen retrieval was performed with boiling 1 mM EDTA, pH 8.0 using a microwave with maintenance at a sub-boiling temperature for 15 min. Antibody concentrations, fluorophore assignments, staining order, TSA incubation times, and imaging settings were determined through optimization experiments using small intestine and tonsil controls and subsequently maintained across all staining batches. Tonsil and small intestine tissues were included in each staining batch as technical controls. Primary antibodies used for mIF staining included CPS1 (#HPA021400, 0.05 mg/ml, Sigma-Aldrich), OLFM4 (#14369, 0.0175 mg/ml, Cell Signaling Technology), ANPEP (#32720, 0.07 mg/ml, Cell Signaling Technology). Detection was performed using species specific HRP conjugated secondary antibodies (all from Cell Signaling Technology) and TSA Plus Fluorescein, Cyanine 5, and Cyanine 3 kit (Akoya Biosciences) for tyramide signal amplification. Order of antibodies and fluorophores was CPS1 (Cyanine 5), OLFM4 (Cyanine 3) and ANPEP (Fluorescein) used sequentially. Stripping of antibodies following signal amplification was performed using boiling 10 mM Sodium Citrate, pH 6.0 in a microwave followed by maintenance at sub-boiling temperature for 10 minutes and cooling on bench top for 30 minutes. Nuclear staining was performed with 2 μg/ml DAPI (Thermo Fisher Scientific).

### Image acquisition

Whole-slide images from mIF were obtained using the Leica Aperio VERSA slide scanner (Leica Biosystems Inc., IL, USA) by the Histology and Light Microscopy Core facility at the Gladstone Institutes, San Francisco, USA. High-resolution fluorescence images were acquired on an LSM 700 confocal microscope (Carl Zeiss) using oil immersion objectives, a 40x EC Plan-Neofluar (NA 1.3) or a 63x Plan-Apochromat (NA 1.4). Brightfield whole-slide images from H&E were obtained using Aperio AT2 whole slide scanner (Leica Biosystems Inc., IL, USA) by the Human Pathology Histology Services core facility at Stanford University.

### Image analysis

WSI image analysis was performed in QuPath version 0.5.2^27^ using steps outlined in the QuPath vignette. Cells were detected using the Watershed algorithm with a requested pixel size of 0.5 µm. Nucleus detection parameters included a background radius of 8 µm with background reconstruction, sigma 0.1 µm, DAPI intensity threshold of 15, and minimum and maximum area of 6 and 400 µm² respectively. Cells were expanded by 6.5 µm from detected nuclei. A composite classifier was generated using mean per-cell fluorescence intensity thresholds for ANPEP, CPS1, and OLFM4. Given the possibility of non-linear signal amplification and saturation, fluorescence intensity was used only for threshold-based classification of marker-positive cells and was not interpreted as a continuous quantitative measure of protein abundance. For each sample, the proportion of cells positive for ANPEP, CPS1, or OLFM4 was calculated as the percentage of all detected cells expressing the respective marker. Differences in marker-positive cell proportions across lesion categories (Normal, GIM-adjacent, GIM-mild, GIM-moderate, and GIM-marked) were assessed using the Kruskal–Wallis test followed by Dunn’s post hoc comparisons with false discovery rate correction. To evaluate trends across metaplastic severity, lesion categories were encoded as an ordinal stage variable (0–4) and analyzed using linear regression, with the proportion of marker-positive cells as the continuous outcome and stage as the predictor.

Confocal microscopy images were processed with Fiji^55^. To generate high-resolution reconstructions of glands, individual image tiles were manually captured with a minimum of 10% overlap and subsequently assembled and blended using the stitching plugin in Fiji.

### Additional analysis

Additional analysis or visualization was conducted using R packages purr (1.1.0), rstatix (0.7.2), tibble (3.3.0), dplyr (1.1.4), tidyr (1.3.1), broom (version 0.7.6), ggplot2 (4.0.0), ggpubr (0.6.1) in R version 4.5.1^56^. Figures were additionally edited in Adobe Illustrator (28.4.1). Pseudocoloring of images was performed in QuPath without any non-linear adjustments.

## AUTHORS’ CONTRIBUTIONS

AS was involved in conception and design of the study, development of methodology, acquisition of data, analysis and interpretation of data and writing of the manuscript. RM was involved in the development of methodology and the acquisition of data. CW was involved in the acquisition of data. BG was involved in the acquisition and interpretation of data. RL performed clinical chart review and managed samples. IAW, SH and JS were involved in the interpretation of data. HPJ and MRA oversaw the conception and design of the study and interpreted the data. RJH oversaw the conception and design of the study, data analysis, interpretation of data and writing of the manuscript.

## ACKNOWLEDGEMENTS

We are grateful to all patients who participated in the study. This work was supported by the US National Institutes of Health grant 5P01CA265772. Schematic figures of small intestine and gastric antral glands in Fig. 5A and B were created using Biorender.com.

## ADDITIONAL FILES

Additional file 1: Supplementary figures S1 – 4. format: PDF

## REFERENCES

1 Bray, F. et al. Global cancer statistics 2022: GLOBOCAN estimates of incidence and mortality worldwide for 36 cancers in 185 countries. CA Cancer J Clin 74, 229–263 (2024). PMID: 38572751. doi:10.3322/caac.21834

2 Correa, P. & Piazuelo, M. B. The gastric precancerous cascade. J Dig Dis 13, 2–9 (2012). PMID: 22188910. doi:10.1111/j.1751-2980.2011.00550.x

3 de Vries, A. C. et al. Gastric cancer risk in patients with premalignant gastric lesions: a nationwide cohort study in the Netherlands. Gastroenterology 134, 945–952 (2008). PMID: 18395075. doi:10.1053/j.gastro.2008.01.071

4 Song, H. et al. Incidence of gastric cancer among patients with gastric precancerous lesions: observational cohort study in a low risk Western population. BMJ 351, h3867 (2015). PMID: 26215280.doi:10.1136/bmj.h3867

5 Li, D. et al. Risks and Predictors of Gastric Adenocarcinoma in Patients with Gastric Intestinal Metaplasia and Dysplasia: A Population-Based Study. Am J Gastroenterol 111, 1104–1113 (2016). PMID: 27185078. doi:10.1038/ajg.2016.188

6 Reddy, K. M., Chang, J. I., Shi, J. M. & Wu, B. U. Risk of Gastric Cancer Among Patients With Intestinal Metaplasia of the Stomach in a US Integrated Health Care System. Clin Gastroenterol Hepatol 14, 1420–1425 (2016). PMID: 27317852. doi:10.1016/j.cgh.2016.05.045

7 Spence, A. D. et al. Adenocarcinoma risk in gastric atrophy and intestinal metaplasia: a systematic review. BMC Gastroenterol 17, 157 (2017). PMID: 29228909. doi:10.1186/s12876-017-0708-4

8 Capelle, L. G. et al. The staging of gastritis with the OLGA system by using intestinal metaplasia as an accurate alternative for atrophic gastritis. Gastrointest Endosc 71, 1150–1158 (2010). PMID: 20381801. doi:10.1016/j.gie.2009.12.029

9 Cho, S. J. et al. Staging of intestinal- and diffuse-type gastric cancers with the OLGA and OLGIM staging systems. Aliment Pharmacol Ther 38, 1292–1302 (2013). PMID: 24134499. doi:10.1111/apt.12515

10 Yue, H., Shan, L. & Bin, L. The significance of OLGA and OLGIM staging systems in the risk assessment of gastric cancer: a systematic review and meta-analysis. Gastric Cancer 21, 579–587 (2018). PMID: 29460004. doi:10.1007/s10120-018-0812-3

11 Latorre, G. et al. Comparison of OLGA and OLGIM as predictors of gastric cancer in a Latin American population: the ECHOS Study. Gut 73, e18 (2024). PMID: 38148138. doi:10.1136/gutjnl-2023-331059

12 Huang, K. K. et al. Spatiotemporal genomic profiling of intestinal metaplasia reveals clonal dynamics of gastric cancer progression. Cancer Cell 41, 2019–2037 e2018 (2023). PMID: 37890493. doi:10.1016/j.ccell.2023.10.004

13 Huang, R. J. et al. A spatial transcriptomic signature of 26 genes resolved at single-cell resolution characterizes high-risk gastric cancer precursors. NPJ Precis Oncol 9, 52 (2025). PMID: 40000871. doi:10.1038/s41698-025-00816-w

14 Mina-Osorio, P. The moonlighting enzyme CD13: old and new functions to target. Trends Mol Med 14, 361–371 (2008). PMID: 18603472. doi:10.1016/j.molmed.2008.06.003

15 Nitzahn, M. & Lipshutz, G. S. CPS1: Looking at an ancient enzyme in a modern light. Mol Genet Metab 131, 289–298 (2020). PMID: 33317798. doi:10.1016/j.ymgme.2020.10.003

16 Wang, X. Y., Chen, S. H., Zhang, Y. N. & Xu, C. F. Olfactomedin-4 in digestive diseases: A mini-review. World J Gastroenterol 24, 1881–1887 (2018). PMID: 29740203. doi:10.3748/wjg.v24.i17.1881

17 Kim, H. et al. Hybrid identity and distinct methylation profiles of incomplete intestinal metaplasia in the stomach. Gut 75, 10–23 (2025). PMID: 40691053. doi:10.1136/gutjnl-2025-335793

18 Tsubosaka, A. et al. Stomach encyclopedia: Combined single-cell and spatial transcriptomics reveal cell diversity and homeostatic regulation of human stomach. Cell Rep 42, 113236 (2023). PMID: 37819756. doi:10.1016/j.celrep.2023.113236

19 Dixon, M. F., Genta, R. M., Yardley, J. H., Correa, P. & the Participants in the International Workshop on the Histopathology of Gastritis, H. Classification and Grading of Gastritis: The Updated Sydney System. The American Journal of Surgical Pathology 20, 1161–1181 (1996). PMID: 00000478-199610000-00001.

20 Patel, A. K., Sethi, N. S. & Park, H. Gastric Cancer: A Review. JAMA 335, 439–450 (2026). PMID: 41499132. doi:10.1001/jama.2025.20034

21 Olsen, J., Kokholm, K., Norén, O. & Sjöström, H. in Cellular Peptidases in Immune Functions and Diseases (eds Siegfried Ansorge & Jürgen Langner) 47–57 (Springer US, 1997).

22 Ghosh, M. et al. Molecular mechanisms regulating CD13-mediated adhesion. Immunology 142, 636–647 (2014). PMID: 24627994. doi:10.1111/imm.12279

23 Uhlen, M. et al. Proteomics. Tissue-based map of the human proteome. Science 347, 1260419 (2015). PMID: 25613900.doi:10.1126/science.1260419

24 Atlas, H. P. <proteinatlas.org> (

25 Vandenberghe-Durr, S., Gilliet, M. & Di Domizio, J. OLFM4 regulates the antimicrobial and DNA binding activity of neutrophil cationic proteins. Cell Rep 43, 114863 (2024). PMID: 39396234. doi:10.1016/j.celrep.2024.114863

26 Kassam, A. F. et al. Olfactomedin 4-Positive Neutrophils Are Upregulated after Hemorrhagic Shock. Am J Respir Cell Mol Biol 64, 216–223 (2021). PMID: 33253592. doi:10.1165/rcmb.2020-0276OC

27 Bankhead, P. et al. QuPath: Open source software for digital pathology image analysis. Sci Rep 7, 16878 (2017). PMID: 29203879. doi:10.1038/s41598-017-17204-5

28 Finkbeiner, S. R. et al. Transcriptome-wide Analysis Reveals Hallmarks of Human Intestine Development and Maturation In Vitro and In Vivo. Stem Cell Reports 4, 1140–1155 (2015). PMID: 26050928. doi:10.1016/j.stemcr.2015.04.010

29 van der Flier, L. G., Haegebarth, A., Stange, D. E., van de Wetering, M. & Clevers, H. OLFM4 is a robust marker for stem cells in human intestine and marks a subset of colorectal cancer cells. Gastroenterology 137, 15–17 (2009). PMID: 19450592. doi:10.1053/j.gastro.2009.05.035

30 Jang, B. G., Lee, B. L. & Kim, W. H. Olfactomedin-related proteins 4 (OLFM4) expression is involved in early gastric carcinogenesis and of prognostic significance in advanced gastric cancer. Virchows Arch 467, 285–294 (2015). PMID: 26070873. doi:10.1007/s00428-015-1793-9

31 Zhang, T. & Tang, X. Refining the diagnostic utility of OLFM4 in gastric cancer precursors: a call for rigorous methodologies. Mol Cancer 23, 161 (2024). PMID: 39118167. doi:10.1186/s12943-024-02077-w

32 Fang, X. et al. Expression profiling of CPS1 in Correa’s cascade and its association with gastric cancer prognosis. Oncol Lett 21, 441 (2021). PMID: 33868479.doi:10.3892/ol.2021.12702

33 Gutierrez-Gonzalez, L. et al. The clonal origins of dysplasia from intestinal metaplasia in the human stomach. Gastroenterology 140, 1251–1260 e1251-1256 (2011). PMID: 21223968. doi:10.1053/j.gastro.2010.12.051

34 Coorens, T. H. H. et al. The somatic mutation landscape of normal gastric epithelium. Nature 640, 418–426 (2025). PMID: 40108450. doi:10.1038/s41586-025-08708-6

35 Alvina, F. B., Chen, T. C., Lim, H. Y. G. & Barker, N. Gastric epithelial stem cells in development, homeostasis and regeneration. Development 150, (2023). PMID: 37746871. doi:10.1242/dev.201494

36 Choi, E. et al. Cell lineage distribution atlas of the human stomach reveals heterogeneous gland populations in the gastric antrum. Gut 63, 1711–1720 (2014). PMID: 24488499. doi:10.1136/gutjnl-2013-305964

37 Leushacke, M., Ng, A., Galle, J., Loeffler, M. & Barker, N. Lgr5(+) gastric stem cells divide symmetrically to effect epithelial homeostasis in the pylorus. Cell Rep 5, 349–356 (2013). PMID: 24209744. doi:10.1016/j.celrep.2013.09.025

38 Barker, N. et al. Lgr5(+ve) stem cells drive self-renewal in the stomach and build long-lived gastric units in vitro. Cell Stem Cell 6, 25–36 (2010). PMID: 20085740. doi:10.1016/j.stem.2009.11.013

39 Hayakawa, Y., Fox, J. G. & Wang, T. C. Isthmus Stem Cells Are the Origins of Metaplasia in the Gastric Corpus. Cell Mol Gastroenterol Hepatol 4, 89–94 (2017). PMID: 28560293. doi:10.1016/j.jcmgh.2017.02.009

40 Han, S. et al. Defining the Identity and Dynamics of Adult Gastric Isthmus Stem Cells. Cell Stem Cell 25, 342–356 e347 (2019). PMID: 31422913. doi:10.1016/j.stem.2019.07.008

41 Willet, S. G. et al. Regenerative proliferation of differentiated cells by mTORC1-dependent paligenosis. EMBO J 37, (2018). PMID: 29467218.doi:10.15252/embj.201798311

42 Goldenring, J. R., Nam, K. T. & Mills, J. C. The origin of pre-neoplastic metaplasia in the stomach: chief cells emerge from the Mist. Exp Cell Res 317, 2759–2764 (2011). PMID: 21907708. doi:10.1016/j.yexcr.2011.08.017

43 Kim, J. et al. Single-cell analysis of gastric pre-cancerous and cancer lesions reveals cell lineage diversity and intratumoral heterogeneity. NPJ Precis Oncol 6, 9 (2022). PMID: 35087207. doi:10.1038/s41698-022-00251-1

44 Wei, H. et al. OLFM4 promotes the progression of intestinal metaplasia through activation of the MYH9/GSK3beta/beta-catenin pathway. Mol Cancer 23, 124 (2024). PMID: 38849840.doi:10.1186/s12943-024-02016-9

45 Nowicki-Osuch, K. et al. Single-Cell RNA Sequencing Unifies Developmental Programs of Esophageal and Gastric Intestinal Metaplasia. Cancer Discov 13, 1346–1363 (2023). PMID: 36929873. doi:10.1158/2159-8290.CD-22-0824

46 Oliver, A. J. et al. Single-cell integration reveals metaplasia in inflammatory gut diseases. Nature 635, 699–707 (2024). PMID: 39567783. doi:10.1038/s41586-024-07571-1

47 Zhang, P. et al. Dissecting the Single-Cell Transcriptome Network Underlying Gastric Premalignant Lesions and Early Gastric Cancer. Cell Rep 27, 1934–1947 e1935 (2019). PMID: 31067475. doi:10.1016/j.celrep.2019.04.052

48 Wen, J. et al. Spatial transcriptomics defines IM-crypt as a premalignant niche with bidirectional divergence in early gastric carcinogenesis. Commun Biol, (2026). PMID: 42098407.doi:10.1038/s42003-026-10179-y

49 Lerch, J. M. et al. Subtyping intestinal metaplasia in patients with chronic atrophic gastritis: an interobserver variability study. Pathology 54, 262–268 (2022). PMID: 35221041. doi:10.1016/j.pathol.2021.12.288

50 Laohawetwanit, T. et al. Histopathologic evaluation of gastric intestinal metaplasia in non-neoplastic biopsy specimens: Accuracy and interobserver reliability among general pathologists and pathology residents. Ann Diagn Pathol 70, 152284 (2024). PMID: 38422806.doi:10.1016/j.anndiagpath.2024.152284

51 Xia, K. et al. GastritisMIL: An interpretable deep learning model for the comprehensive histological assessment of chronic gastritis. Patterns (N Y) 6, 101286 (2025). PMID: 40843346. doi:10.1016/j.patter.2025.101286

52 Solmaz, O. A. & Tasci, B. ViSwNeXtNet Deep Patch-Wise Ensemble of Vision Transformers and ConvNeXt for Robust Binary Histopathology Classification. Diagnostics (Basel) 15, (2025). PMID: 40564828. doi:10.3390/diagnostics15121507

53 Cano, F. et al. Towards deep-learning based detection and quantification of intestinal metaplasia on digitized gastric biopsies: a multi-expert comparative study. Sci Rep, (2026). PMID: 41741481.doi:10.1038/s41598-025-32737-w

54 Sathe, A. et al. GITR and TIGIT immunotherapy provokes divergent multicellular responses in the tumor microenvironment of gastrointestinal cancers. Genome Med 15, 100 (2023). PMID: 38008725. doi:10.1186/s13073-023-01259-3

55 Schindelin, J. et al. Fiji: an open-source platform for biological-image analysis. Nat Methods 9, 676–682 (2012). PMID: 22743772. doi:10.1038/nmeth.2019

56 A language and environment for statistical computing (R Foundation for Statistical Computing, Vienna, Austria 2022).

