## Supplementary figures for "Basal gland localization and focal distribution of OLFM4-expressing cells in increasing severity of gastric intestinal metaplasia"

**INSTITUTIONS**

[1] Division of Oncology, Department of Medicine, Stanford University School of Medicine, Stanford, CA, USA

[2] Department of Microbiology and Immunology, Stanford University, Stanford, CA, USA.

[3] Department of Pediatrics, Stanford University, Stanford, CA, 94305

[4] Division of Gastroenterology and Hepatology, Department of Medicine, Stanford School of Medicine, Stanford, CA, USA

[5] Quantitative Sciences Unit, Department of Medicine, Stanford University School of Medicine, Stanford, CA, USA

[6] Department of Neurosurgery, Stanford University School of Medicine, Stanford, CA, USA

[7] Department of Pathology, Stanford University, Stanford, CA, USA

**CORRESPONDING AUTHORS**

Anuja Sathe

Mailing address: CCSR 1120, 269 Campus Drive, Stanford, CA-94305, USA

Robert J. Huang

Mailing address: 300 Pasteur Drive, Alway Building M211, Stanford, CA 94035, USA

Conflict of interest statement: The authors declare no potential conflicts of interest.

**SUPPLEMENTARY FIGURE LEGENDS**

**Supplementary Figure 1. mIF staining of small intestine and tonsil tissues.** (A) Multiplex immunofluorescence whole slide image from small intestine (scale bar: 100 μm) (top left). Rectangle with white box in whole slide image indicates area that was imaged using confocal microscopy in all other images (scale bar: 50 μm). All images are pseudocolored for DAPI (blue), ANPEP (green), CPS1 (red) and OLFM4 (yellow). (B) Multiplex immunofluorescence whole slide image from tonsil (scale bar: 100 μm) pseudocolored for DAPI (blue), ANPEP (green), CPS1 (red) and OLFM4 (yellow).

**Supplementary Figure 2. mIF staining of GIM.** Confocal image from sample P68 displaying respective proteins. Scale bars: 20 μm. Images are pseudocolored for DAPI (blue), ANPEP (green), CPS1 (red) and OLFM4 (yellow)

**Supplementary Figure 3. Expression of ANPEP, CPS1 and OLFM4 in GIM.** (A-C) Histogram distribution of mean intensity per cell for respective proteins. Dotted line indicates threshold used in the image classifier. (D-F) Linear regression analysis testing the association between the proportion of marker-positive cells and lesion severity. Lesion categories were encoded as an ordinal stage variable. Each point represents one sample; the line indicates the fitted regression with 95% confidence interval, with regression coefficient (β), coefficient of determination ($R^2$), and p-value. (G) Proportion of cells positive for the expression of respective proteins in dysplasia and cancer samples. Each point represents an individual sample, and boxplots show the median and interquartile range.

**Supplementary Figure 4. Spatial distribution of ANPEP, CPS1 and OLFM4 in gastric tissues.** Multiplex immunofluorescence whole slide images of respective samples across lesion categories. Fluorescence images are pseudocolored for DAPI (blue), ANPEP (green), CPS1 (red) and OLFM4 (yellow). Scale bars: 50 μm.

**Supplementary figure 1**

**A**

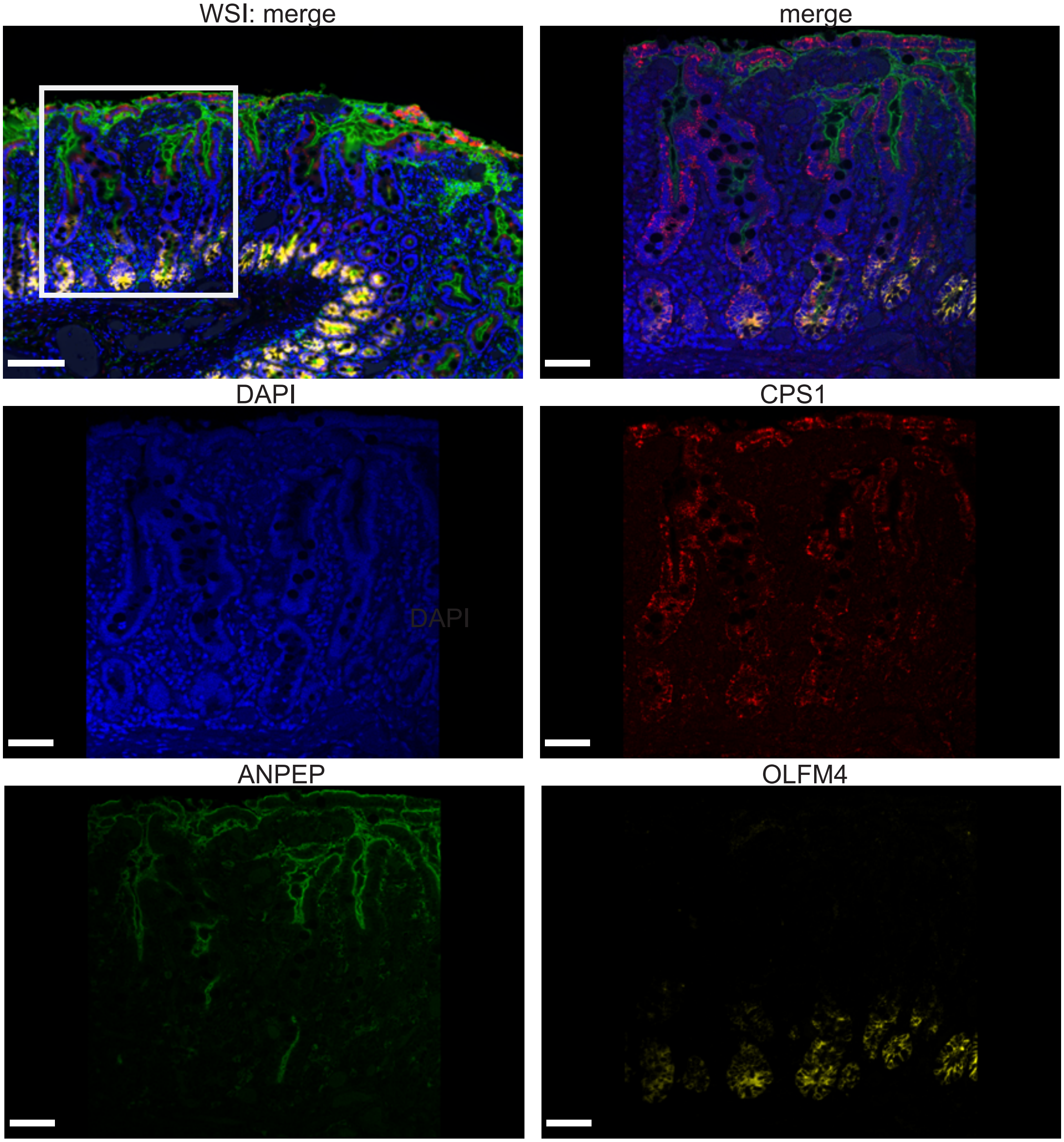


**B**

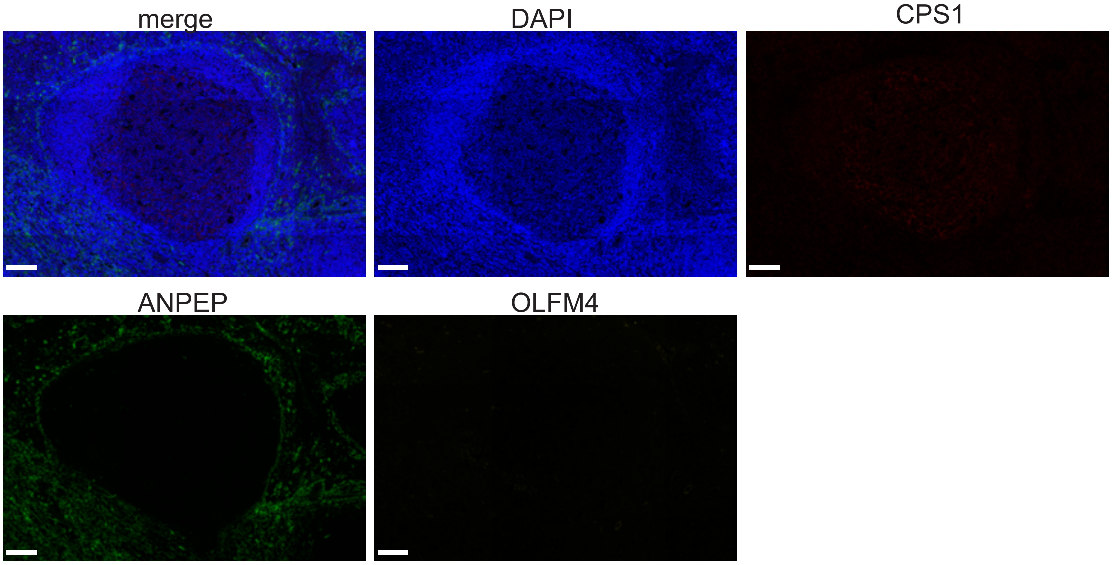

**Supplementary figure 2**

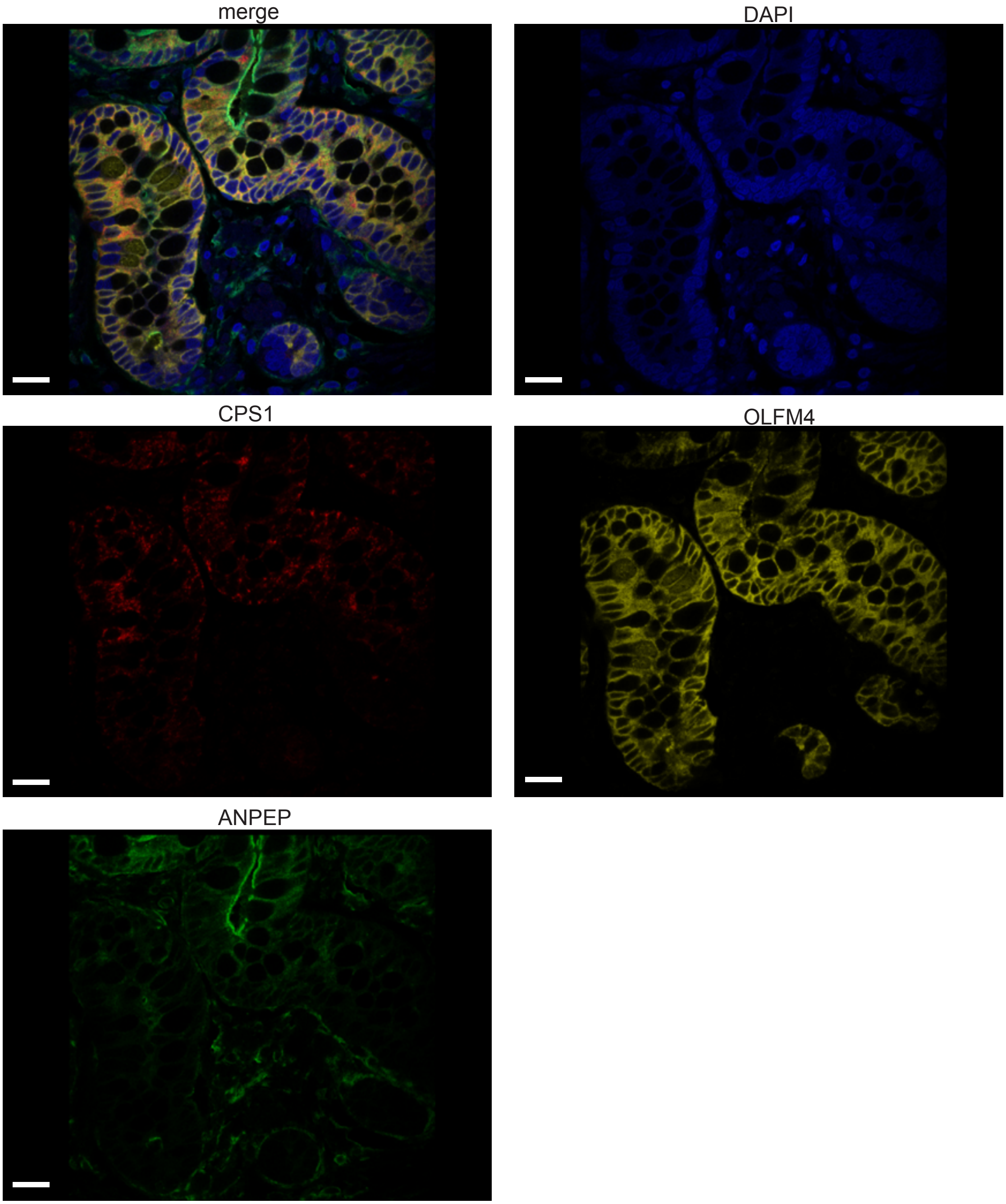

### Supplementary figure 3

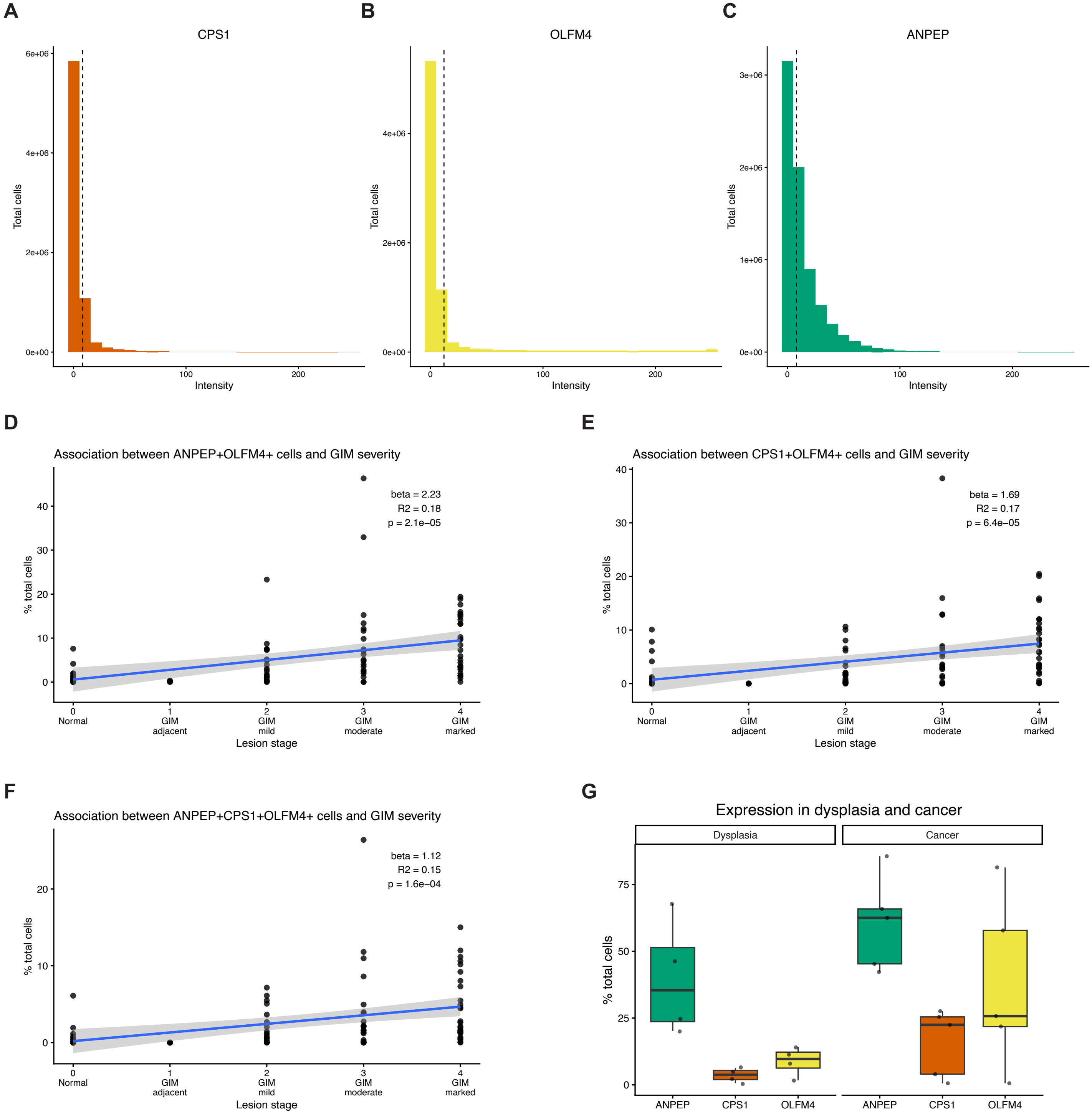

**Supplementary figure 4**

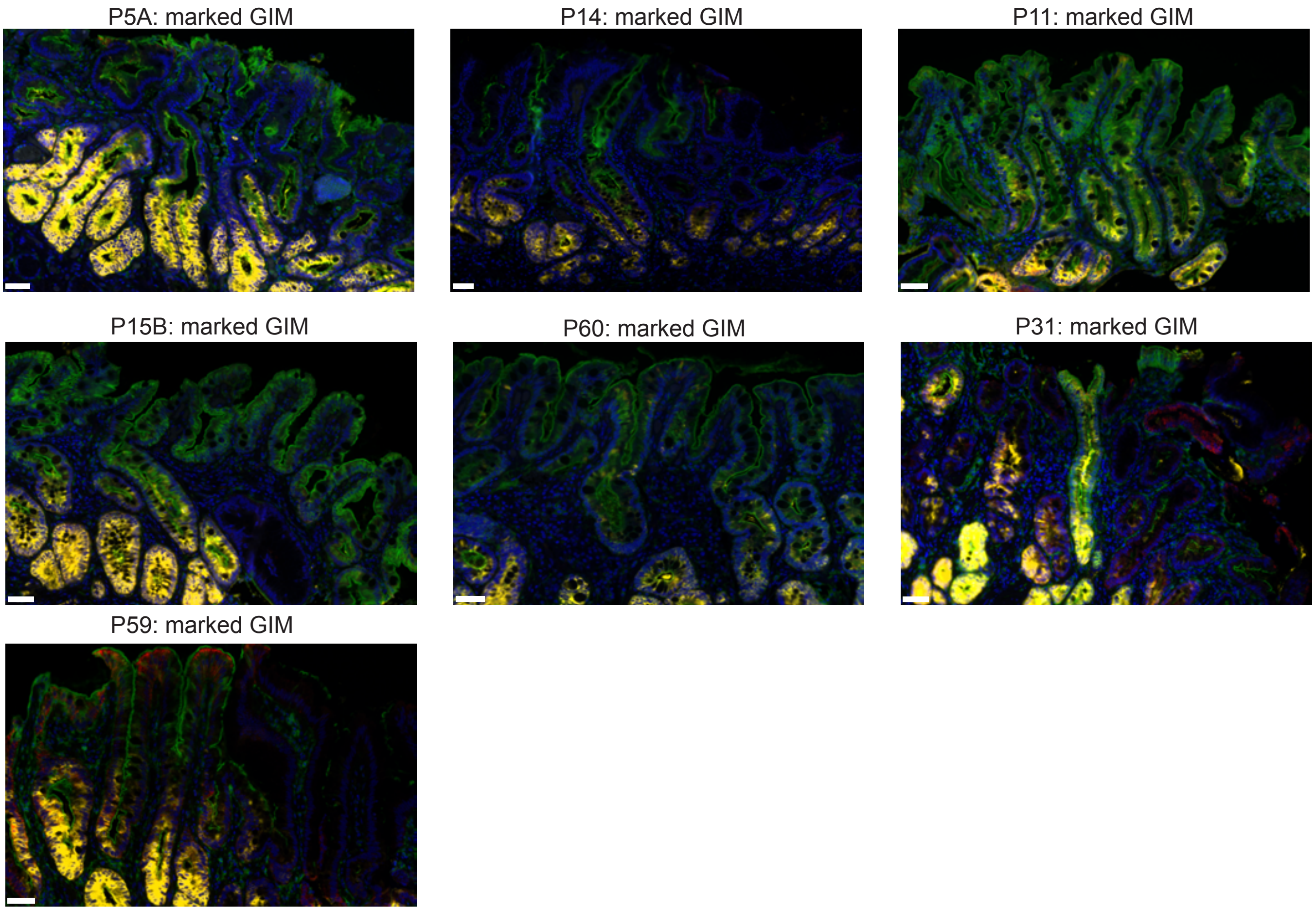
